# Late Pleistocene polar bear genomes reveal the timing of allele fixation in key genes associated with Arctic adaptation

**DOI:** 10.1101/2023.11.30.569368

**Authors:** Yulin Sun, Eline D. Lorenzen, Michael V. Westbury

## Abstract

The polar bear (*Ursus maritimus*) occupies a relatively narrow ecological niche, with many characteristics adapted for cold temperatures, movement across snow, ice and open water, and for consuming highly lipid-dense prey species. The divergence of polar bears from brown bears (*Ursus arctos*) and their adaptation to their Arctic lifestyle is a well-known example of rapid evolution. Previous research investigating whole genomes uncovered genomic regions containing an array of key genes highly differentiated between polar and brown bears, many of which were linked to the novel Arctic environment. Further research suggested fixed alleles in these genes arose from selection on both standing variation and *de novo* mutations in the evolution of polar bears. Here, we reevaluate these findings by incorporating more genomic data from previously unavailable polar and brown bear populations, and assess the timing of allele fixation by utilising the genomes of two Late Pleistocene polar bears (aged 130-100,000 years old and 100-70,000 years old). Contrary to previous results, we found no evidence for alleles fixed in all polar bears within these key genes arising from *de novo* mutation. Most alleles fixed in modern polar bears were also fixed in the Late Pleistocene bears, suggesting selection occurred prior to 70,000 years ago. However, some sites fixed in modern polar bears were not fixed in the Late Pleistocene bears, including at sites within APOB, LYST, and TTN. The functions of these three genes are associated with the cardiovascular functions, metabolism, and pigmentation, suggesting that selection may have acted on different loci at different times.

## Introduction

The polar bear (*Ursus maritimus*) is uniquely adapted to the extreme conditions of life in the High Arctic and spends most of its life out on sea ice. In cold Arctic climates, energy is in high demand. As a result, the polar bear feeds on a lipid-rich diet throughout its life [1]. Polar bears are most closely related to the brown bear (*Ursus arctos*), a widely distributed omnivore found in a variety of habitats across the Holarctic [2]. The two species differ fundamentally in their ecology, behaviour, and morphology, reflecting adaptations to different ecological niches. Polar bears diverged from brown bears relatively recently – within the past 500,000 years [3].

A previous study reported key genes that showed a signal of strong positive selection in the polar bear lineage [3]. These key genes may have played significant roles in the ability of polar bears to rapidly adapt to their new Arctic environment. They included APOB, LYST, and TTN, which are related to cardiovascular functions (APOB, TTN), metabolism (APOB, LYST), and pigmentation (APOB, LYST).

Further research utilising all polar and brown bear genomes available at the time investigated whether alleles fixed in those same key genes in polar bears were derived via selection on standing variation, or on *de novo* mutations unique to the polar bear lineage. Results found evidence of both, suggesting variation already present in the ancestral gene pool and *de novo* mutations played an important role in the evolution of polar bears [4].

Here, we build on this work by incorporating additional, recently published polar bear and brown bear genomes from previously unstudied populations [5–7]to more precisely characterise whether the alleles in the previously identified key genes originate from ancestral standing variation or *de novo* mutation in the polar bear lineage. By incorporating a larger dataset, [5–7] we minimise the possibility of missing data and/or population structure having influenced previous inferences. Furthermore, we incorporate genomic data from two Late Pleistocene polar bears aged 130-100,000 years old (‘Poolepynten’, Svalbard) and 100-70,000 years old (‘Bruno’, Alaska) [2, 8], to investigate the timing of fixation. Establishing a reliable time frame for when the alleles in the key genes showing a signal of strong selection in polar bears become fixed can improve our understanding of what evolutionary processes drove speciation, and the rates in which novel adaptations to extreme environments can arise.

## Results

### Ancient DNA damage

We observed high levels of C-T substitution towards the ends of the reads and G-A on the reverse complement in both Late Pleistocene samples (Supplementary Fig 1). Bruno, the Late Pleistocene polar bear individual from Alaska, displayed less damage with ∼5% of the sites at the read ends experiencing ancient DNA (aDNA) damage. Poolepynten, the Late Pleistocene polar bear individual from Svalbard, had more damage with ∼20% of the sites at the read ends showing aDNA damage patterns.

### Principal Component Analysis

To investigate whether the enlarged dataset may have uncovered gene flow between polar and brown bears in the genomic regions containing the key genes, we performed independent principal component analyses for each gene plus their 50kb flanking regions. In all eleven principal component analyses we observed clear differentiation between polar and brown bear individuals, suggesting no interspecific admixture at these loci (Supplementary Figs S2-S12).

### Fixed derived alleles in polar bears

When comparing genotype calls between all modern polar and brown bears, we found no evidence for any of the alleles in the eleven focus genes to have arisen by *de novo* mutations in the polar bear lineage. That is, we found no sites that were fixed for the polar bear reference genome allele in polar bear and fixed for the alternative allele in the brown bear, giant panda (*Ailuropoda melanoleuca*), and spectacled bear (*Tremarctos ornatus*) (Fig 2, Supplementary table S1). Four genes (CUL7, FCGBP, LAMC3, XIRP1) contained no sites that were fixed for the derived allele in the polar bear (Supplementary table S1).

**Figure 1:**
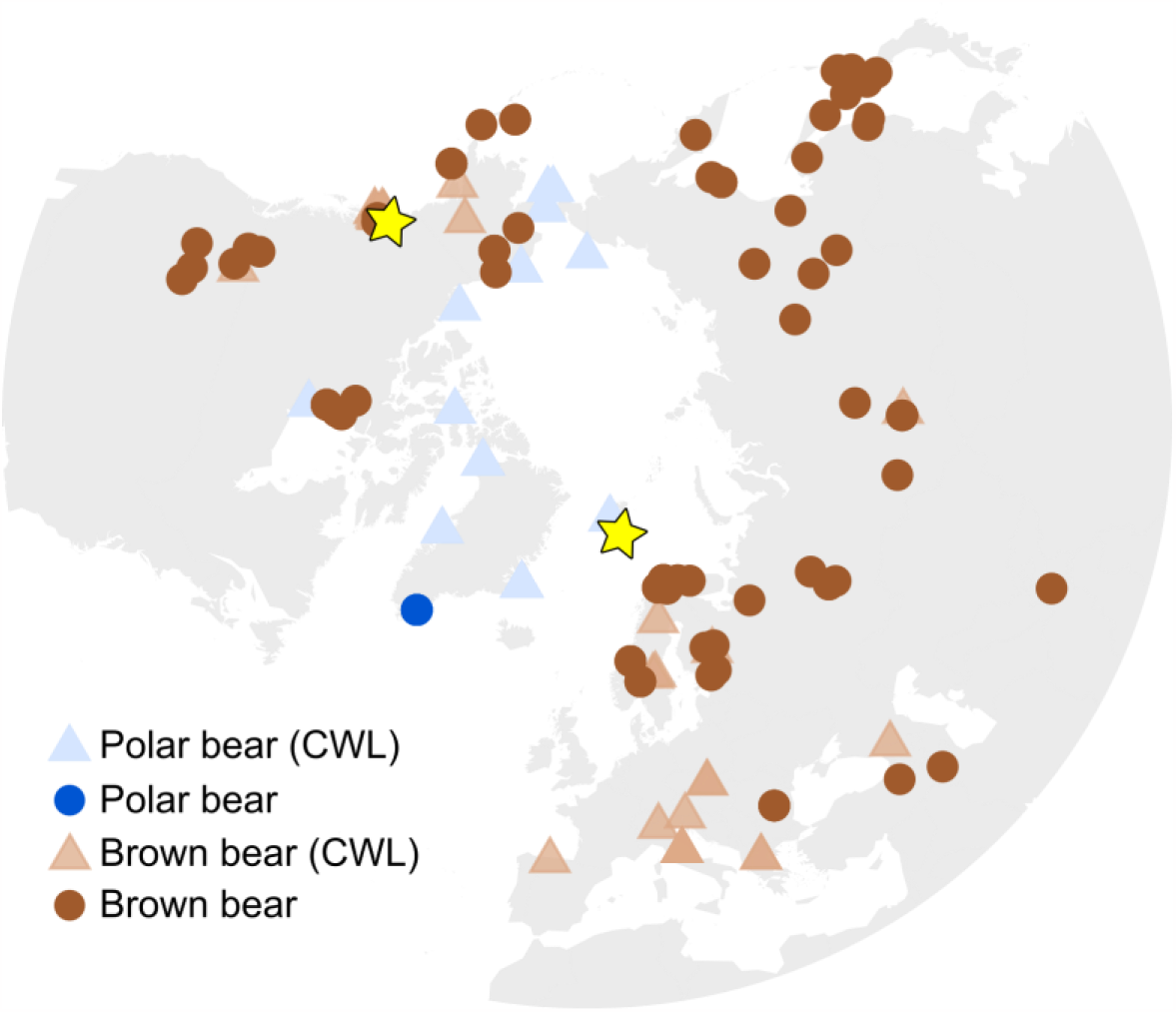
Geographic localities of the polar and brown bears included in this study. CWL shows the bears that were used in the study by Castruita, Westbury, and Lorenzen. Stars indicate the Late Pleistocene polar bears.

**Figure 2:**
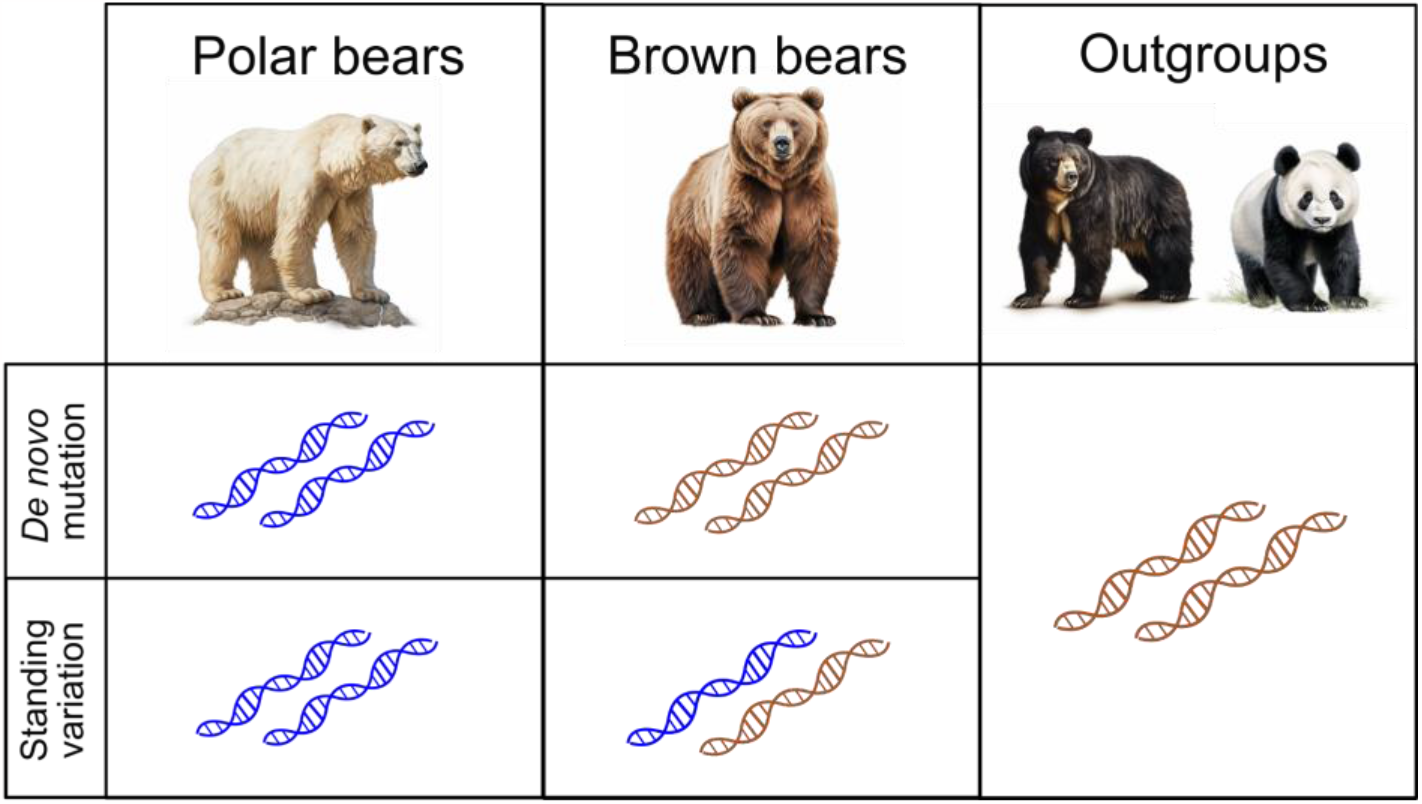
Schematic overview of how we determined whether alleles fixed in all modern polar bears arose through *de novo* mutation in the polar bear lineage or whether they were present as standing variation in the ancestral brown/polar bear gene pool. Images generated using the AI tool Midjourney.

### Timing of allele fixation in polar bears

All seven genes that remained after filtering had sites that showed signs of the derived allele being fixed prior to the age of the Late Pleistocene individuals; the Late Pleistocene individuals were also fixed for the derived allele (Fig 3, Table 1, Supplementary table S1). Only APOB, LYST and TTN contained sites showing signs of the derived allele becoming fixed after the age of the Late Pleistocene individuals; at least one of the Late Pleistocene individuals also contained the ancestral allele (Table 1, Supplementary table S1).

**Table 1.**
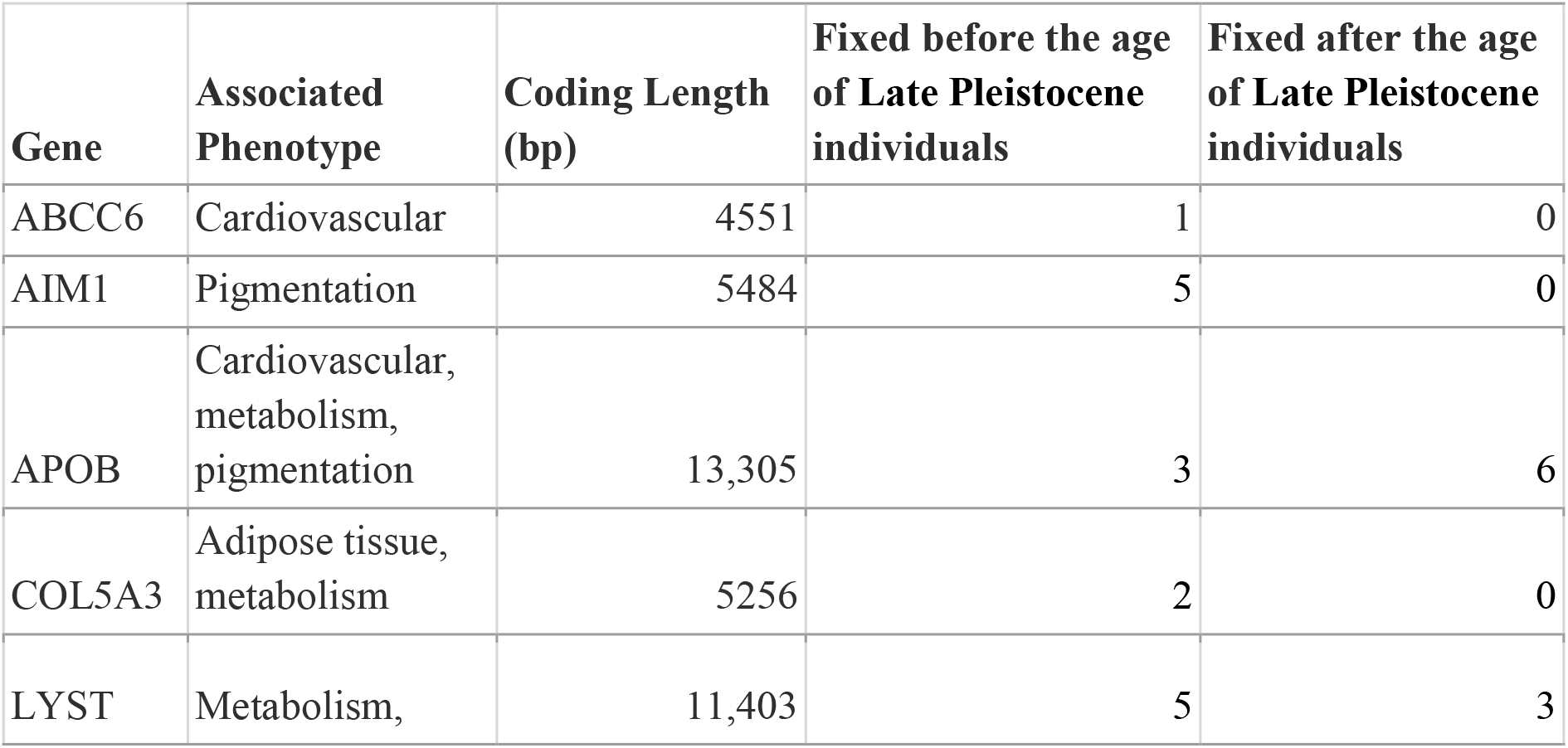
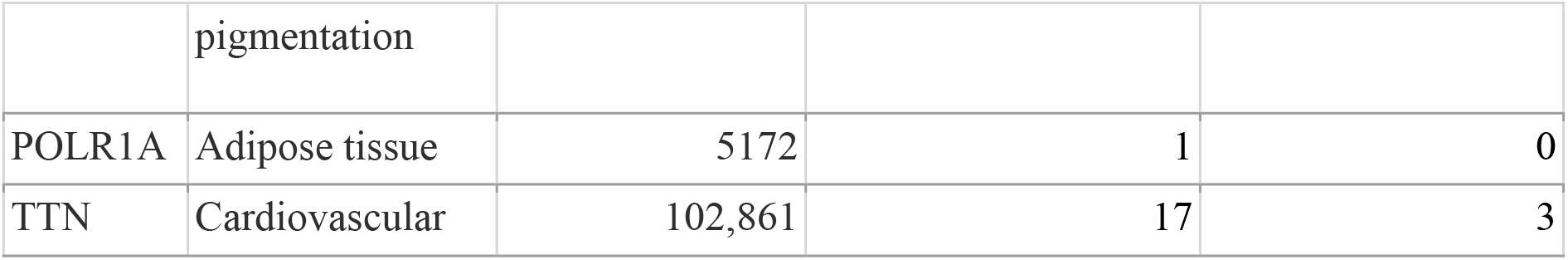
Timing of fixation of alleles in genes previously identified to show a signature of strong selection in the polar bear lineage. Genes with no fixed alleles in polar bears after filtering are not shown.

| Gene | Associated Phenotype | Coding Length (bp) | Fixed before the age of Late Pleistocene individuals | Fixed after the age of Late Pleistocene individuals |
| --- | --- | --- | --- | --- |
| ABCC6 | Cardiovascular | 4551 | 1 | 0 |
| AIM1 | Pigmentation | 5484 | 5 | 0 |
| APOB | Cardiovascular, metabolism, pigmentation | 13,305 | 3 | 6 |
| COL5A3 | Adipose tissue, metabolism | 5256 | 2 | 0 |
| LYST | Metabolism, pigmentation | 11,403 | 5 | 3 |
| POLR1A | Adipose tissue | 5172 | 1 | 0 |
| TTN | Cardiovascular | 102,861 | 17 | 3 |

**Figure 3:**
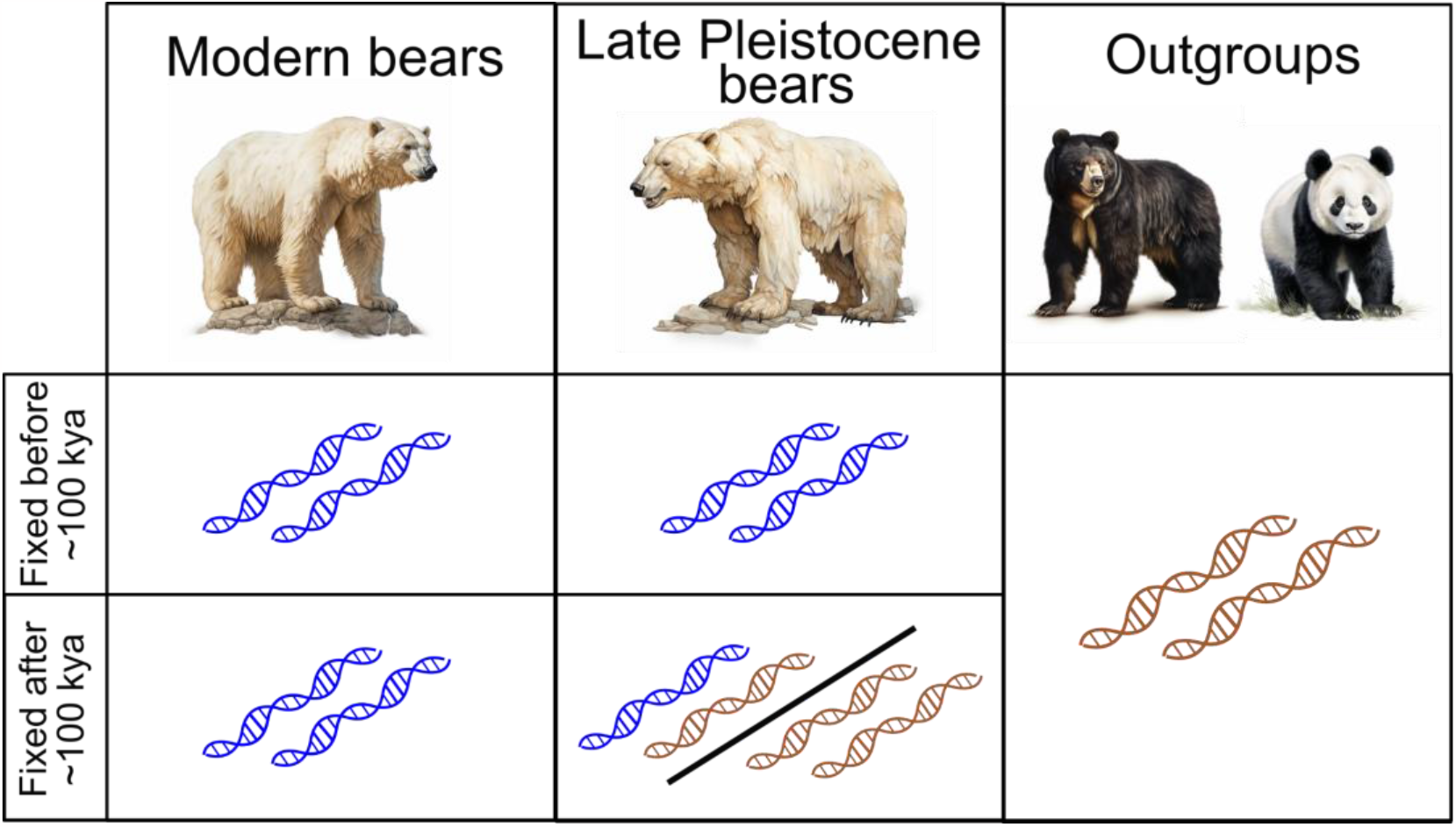
Schematic overview of how we determined whether an allele was fixed in the polar bear lineage before or after the age of the Late Pleistocene polar bears. Images generated using the AI tool Midjourney.

### Assessing heterozygous base call reliability

To avoid biases due to aDNA damage, we set the threshold of minor allele frequency of 30% to decide whether a heterozygous base call in the Late Pleistocene individuals was a false positive. We found 14 heterozygous sites in the Late Pleistocene individuals (Supplementary table S2), two of which may be false positives in the TTN gene (minor allele frequencies of 25% and 29%). However, through visualisation of the mapped reads we found that the minor alleles always occurred within the read as opposed to the end. As aDNA damage occurs mostly towards the ends of the reads, and they were close to the 30% cutoff, we designated these as true heterozygous sites.

## Discussion

By utilising genomes from two Late Pleistocene polar bears aged 110-70 thousand years old and 130-110 thousand years old, we were able to make inferences about the timing in which alleles found in genes proposed to have driven the differentiation between polar and brown bears [3], become fixed in the polar bear lineage. We found 34 sites that were fixed in all modern and the two Late Pleistocene polar bears suggesting fixation of these alleles occurred prior to the ages of the Late Pleistocene bears. We found 13 sites containing alleles that were fixed in all modern polar bears but not in the Late Pleistocene bears, suggesting their fixation occurred after the age of the Late Pleistocene bears.

Our result of more sites becoming fixed prior to the age of the Late Pleistocene bears is congruent with what we know about the Late Pleistocene Poolepynten bear. Based on comparisons of morphometric measurements of its well-preserved mandible with modern and fossil polar bears, as well as brown bears, it was shown to fall within the range of modern polar bears [9]. Stable isotopes also revealed it to be subsisting on a marine diet. Therefore, we can assume this individual already possessed key traits and was adapted to the Arctic environment.

The 13 derived alleles that became fixed after the ages of the Late Pleistocene bears (minimally ∼70,000 years ago), were found in only three key genes: APOB, LYST, and TTN. Although it is difficult to identify the determinant allele for a phenotype, this result suggests these alleles may not have been imperative in first enabling the polar bear to evolve to an Arctic existence. However, the functions of these three genes are associated with the cardiovascular functions, metabolism, and pigmentation, suggesting the genes themselves played a role in polar bears’ Arctic adaptation.

The gene showing the most alleles with signs of becoming fixed after the ages of the Late Pleistocene polar bears was APOB, as there are relatively more derived alleles that are not fixed in the Late Pleistocene bears. The gene APOB codes for apolipoprotein B (apoB), which is associated with the cardiovascular system [10]. It has been suggested that selection on the APOB gene may have played a role in the novel adaptation of polar bears to a lipid-rich diet, and increased the efficacy of cholesterol clearance from the blood [3, 11]. The feeding ecology of Poolepynten was shown to fall within the range of present-day polar bears, who prey mainly on ringed seals and bearded seals [9]. Therefore, we can assume that the ability to process a lipid-rich diet was required more than 70,000 years ago, suggesting selection cannot have driven this phenotype within the last 70,000 years. This lends to the idea that the variants we discuss here may not have been pivotal in the early adaptation, but may have been driven by increased selective pressures during the last glacial period. Other genes showing a strong signal of selection in the polar bear lineage, such as ABCC6 and COL5A3, also have functions related to the cardiovascular system and metabolism [4]. Only they have alleles fixed in the Late Pleistocene bears, and they may therefore have played a key role in driving the early adaptation of polar bears to a lipid-rich diet.

Similar to *APOB, TTN* is associated with the cardiovascular system. TTN encodes Titin, an abundant protein of striated muscle, which includes cardiac muscle tissue; mutations in *TTN* are linked with human cardiac physiology [12]. The gene LYST is associated with pigmentation [13]. Pigmentation is a phenotype that is not preserved in the fossil record, therefore it is hard to know when the white phenotype of polar bears evolved. As both of these genes contain more alleles that were fixed in the Late Pleistocene polar bears, the alleles in these genes not fixed in the Late Pleistocene bears may have either been driven to fixation in the modern polar bears by selection in the last ∼70,000 years or by linkage disequilibrium, rather than selection.

Contrary to the previous study[4], we did not find any signs of *de novo* mutation in any of the eleven genes investigated. At all sites, both the ancestral and the derived alleles were found in brown bears, suggesting both were also present in the ancestral brown/polar bear gene pool. The addition of substantially more brown bear individuals (Fig 1) to this study decreased the chances of allelic drop out. In the case of the polar bear, which rapidly adapted to its novel Arctic environment, the lack of *de novo* mutations is perhaps not surprising. While both standing variation and *de novo* mutations provide the raw material for evolution, standing variation is already in the gene pool for selection to act upon, allowing for immediate use in adaptation. *De novo* mutations arise randomly, segregate at an initially low frequency, and therefore need more time to reach fixation under the same selective pressure [14]. The above suggests that in polar bear populations, no matter when the selection occurred, standing variation was key to the ability to survive the Arctic environment.

Our study provides novel evidence of the timing and modes of selection. However, palaeogenomic data are only available from two Late Pleistocene polar bear individuals. Consequently, inferences regarding the timing of allele fixation must be interpreted with caution. Specifically, the fixation of a given allele in only two individuals cannot be confidently extrapolated to the wider population which existed during the Late Pleistocene, although they are geographically disparate and therefore may represent a wide breadth of the polar bear ancestry of the time. However, additional ancient data, which may become available in the future, may enable us to further pinpoint the timing of these crucial adaptations, which have enabled polar bears to inhabit one of the coldest environments on Earth

## Methods

### Pleistocene polar bear individuals

Available data from two Pleistocene polar bears were included in the study. Genomic data of Bruno (110-70,000 years old) was previously generated from the skull of a juvenile polar bear sample that was found in 2009 on the coast of the Beaufort Sea, near Point McLeod in Arctic Alaska [2]. Genomic data from Poolepynten (130-110,000 years old) was previously extracted from a left mandible, which was found in Svalbard. Age determination with infrared-stimulated luminescence suggested that it is probably the oldest polar bear fossil discovered [8, 15].

### Modern individuals

Following the previous study by Castruita, Westbury, and Lorenzen [4], our analysis included the data set from Liu et al of 89 genomes [3] (79 polar bear, 10 brown bear) and the 30 polar bear and 23 brown bear genomes published elsewhere [16–20]. We obtained the mapped files from the Castruita, Westbury, and Lorenzen publication which utilised raw reads from NCBI (Bioproject IDs: PRJNA169236, PRJNA196978, PRJNA210951, PRJNA271471, PRJNA395974, and PRJEB27491).

New to the present study, we incorporated newly available genomic data from populations of polar bears in Southeast Greenland (n = 10) [5], and brown bears from Hokkaido, Japan (n = 6)[7] and across their Holarctic distribution (n=96) [6]. We downloaded the SRA files from NCBI from the Bioproject IDs: PRJNA669153, PRJDB11280, and PRJNA913591. Information on the newly incorporated individuals can be found in supplementary table S3.

### Raw data processing

For the 142 individuals from Castruita, Westbury, and Lorenzen [4], raw sequencing reads were previously processed with the PALEOMIX [21] pipeline. Internally, adapter sequences, stretches of Ns, and low-quality bases were trimmed and filtered with AdapterRemovalv2 [22] using default parameters. BWA v0.7.15 [23] aln was used to map the cleaned reads to the pseudo-chromosomal polar bear genome (Genbank accession: GCA_000687225.1) from Liu et al [3], with default parameters. Reads with mapping quality of less than 30 were filtered using SAMtools v1.6 [24]. Duplicates were removed with picard v2.6.0 [25]. Possible paralogs were filtered using SAMtools. Finally, local realignment around indels was performed using GATK (v 3.3) [26].

For the 112 newly incorporated individuals, we trimmed adapter sequences and merged overlapping read pairs with Fastp v0.23.2 [27]. We mapped the cleaned reads to the same pseudochromosome polar bear genome with BWA v0.7.15 [28] aln with the seed disabled (-l 690). We used SAMtools v1.6 [24] to filter the reads with mapping quality of less than 30 and remove duplicates. We assessed aDNA damage in the two Late Pleistocene polar bear individuals and adjusted base quality scores around damage using Mapdamage2 (--rescale) [29, 30]. To determine the ancestral allele, we included single individual representatives of the spectacled bear and the giant panda (Bioprojects PRJNA472085 and PRJNA38683). We mapped the reads to the same pseudochromosome polar bear genome following the same approach as the 112 newly incorporated modern individuals.

### Population structure and admixture

To investigate whether admixture or incomplete lineage sorting (ILS) may be present between polar bears and brown bears at the eleven genes of interest, we performed independent principal component analyses (PCAs) for each gene including the 50 kb regions upstream and downstream of the gene. We used a genotype likelihood approach to construct the PCAs: input genotype likelihood files were constructed using ANGSD v0.929 [31], with the SAMtools genotype likelihood algorithm (–GL 1), and specifying the following parameters: remove reads that have multiple mapping best hits (–uniqueonly), remove reads with a flag above 255/secondary hits (–remove_bads), include only read pairs with both mates mapping correctly (–only_proper_pairs), adjust mapQ for reads with excessive mismatches (–C 50), adjust quality scores around indels (–baq 1), a minimum mapping quality of 20 (–minMapQ 20), a minimum base quality of 20 (–minQ 20), determine the major allele based on the maximum likelihood approach (-doMajorMinor 1), calculate allele frequencies assuming a fixed major allele and an unknown minor allele (-doMaf 2), generate beagle input file (-doGlf 2), discard sites where there is no data in at least 95% of the individuals (–minInd), skip tri-allelic sites (–skipTriallelic), and remove SNP sites with a p-value larger than 1e– 6 (–SNP_pval 1e-6). The ANGSD output beagle file was run through PCAngsd v0.95 [32] to generate a covariance matrix.

### Genotype calling

We investigated eleven of the genes previously inferred using population genomics and demographic modelling to have the strongest signals of positive selection in the polar bear [3]. These included ABCC6, AIM1, APOB, COL5A3, CUL7, FCGBP, LAMC3, LYST, POLR1A, TTN, and XIRP1. We excluded EDH3 as in, due to potential for admixture between polar and brown bears in the genomic region containing the gene[4].

We called genotypes using ANGSD v0.921 [31] following the approach of [4]. We used the SAMtools genotype likelihood algorithm (-GL 1), remove reads that have multiple mapping best hits (-unique_only 1), remove reads with a flag above 255/secondary hits (-remove_bads 1), adjust quality scores around indels (-baq 1), a minimum mapping quality of 20 (-minMapQ 20), a minimum base quality of 20 (-minQ 20), write major and minor alleles and the genotype directly (-doGeno 5), estimate the posterior genotype probability based on the allele frequency as a prior (-doPost 1), use the reference allele as the major allele (-doMajorMinor 4), and calculate allele frequencies assuming a fixed major allele and an unknown minor allele (-doMaf 2). In order to decrease biases that could arise when calling heterozygous alleles from the low-coverage genomes, we only called genotypes from individuals that had at least 4x coverage at the site of interest (-geno_minDepth 4). We only included biallelic sites where each allele led to a different amino acid (non-synonymous differences).

To determine which allele was the ancestral allele, we used the outgroup spectacled bear and giant panda sequences. If the allele fixed in all polar bears was found in either of these individuals, we removed that site from further consideration.

We further investigated for false positive heterozygous sites that may have arisen due to aDNA damage in the Late Pleistocene individuals. We investigated the read count for each of the four bases at each site of interest, focusing on heterozygous sites which might be caused by aDNA damage (C-T and G-A). Read counts were generated in ANGSD using the - dumpcount parameter. We calculated the proportion of the minor base of each heterozygous site and only if the ratio is more than 30%, would we assume that this site is heterozygous and not a false positive.

## Declarations

### Ethics approval and consent to participate

Not applicable

### Consent for publication

Not applicable

### Availability of data and materials

All polar and brown bear short read data can be found under the following NCBI Bioproject IDs: PRJNA169236, PRJNA196978, PRJNA210951, PRJNA271471, PRJNA395974, PRJEB27491, PRJNA669153, PRJDB11280, and PRJNA913591. The polar bear genome used as the mapping reference can be found under the Genbank accession: GCA_000687225.1. The spectacled bears and giant panda short read data found under the following NCBI Bioproject IDs PRJNA472085 and PRJNA38683.

### Author’s contributions

E.D.L and M.V.W conceptualised the study. Y.S performed the computational analyses. Y.S and M.V.W interpreted the results. Y.S and M.V.W wrote the initial draft of the manuscript. All authors read and approved the final manuscript.

## Acknowledgements

Not applicable

