## Supplementary information for "Late Pleistocene polar bear genomes reveal the timing of allele fixation in key genes associated with Arctic adaptation"

**Supplementary tables**

**Table S1.** Gene location, nucleotide information, allele counts, and amino acid information for the genes of interest. 'R' indicates the reference allele count. 'A' indicates the alternative allele count. Amino acid information taken from Castruita, Westbury, and Lorenzen 2020.

**Table S2.** Heterozygous sites (marked by yellow), base counts and the proportion of the minor allele at the highlighted heterozygous site. The base count is only shown for the heterozygous individual.

**Table S3**: Geography, genome-wide mean coverage, and reference for the newly included polar and brown bear genomes, including the two ancient polar bears.

### Supplementary figures

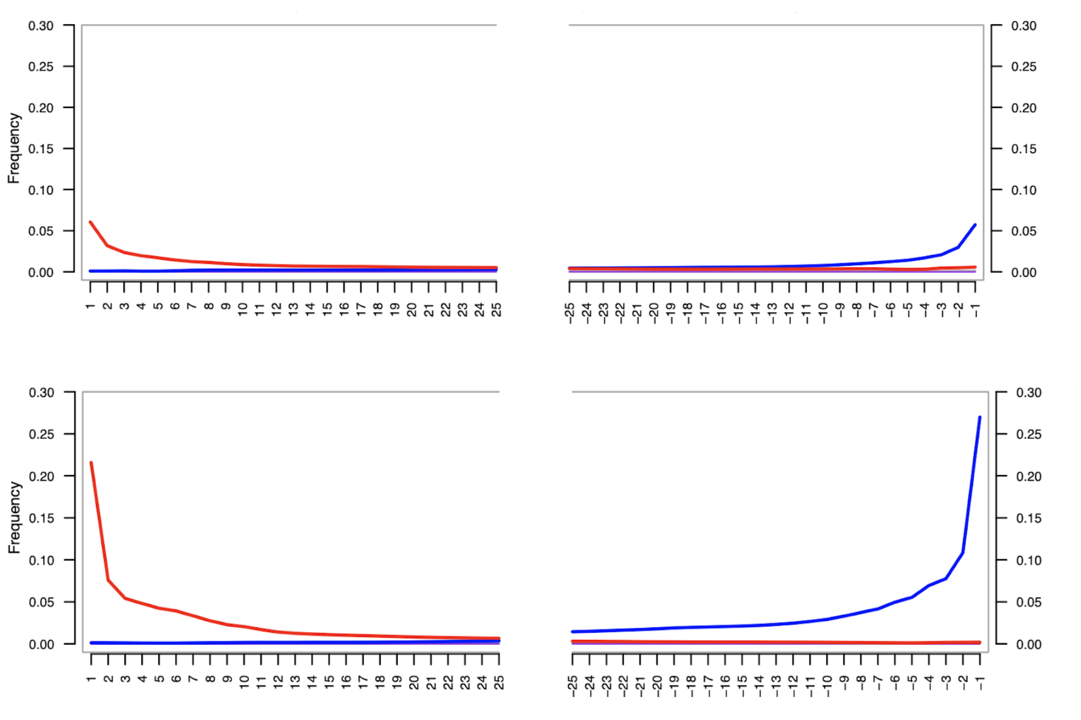


**Figure S1**. Mapdamage results. Upper plot shows the damage patterns plot for Bruno, while the lower plot shows the damage patterns for Poolepynten. Red indicates C to T transitions, blue indicates G to A transitions. Y -axis denotes the proportions of sites containing a nucleotide change from the reference sequence. X-axis represents position from 5' (left) and 3' (right) read end.

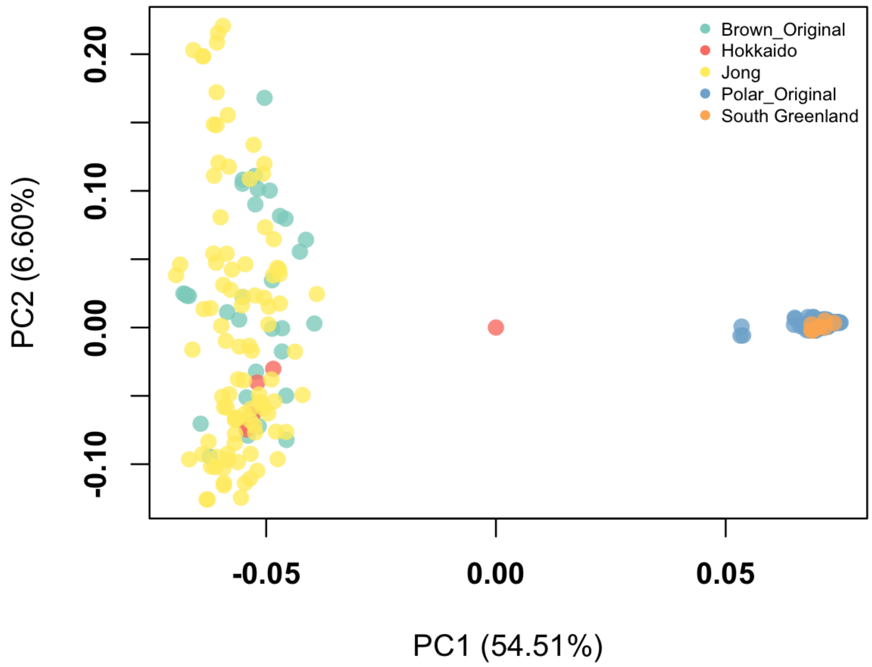


**Figure S2**. Principal component analysis of gene AIM1 and the 50kb flanking regions using all modern individuals included in this study. 'Polar_Original' and 'Brown_Original' refer to the samples from Castruita, Westbury, and Lorenzen 2020. 'South Greenland', 'Hokkaido', represent new polar bear datasets from Laidre et al and Endo et al respectively. 'Jong' shows the Holarctic brown bear dataset from de Jong et al.

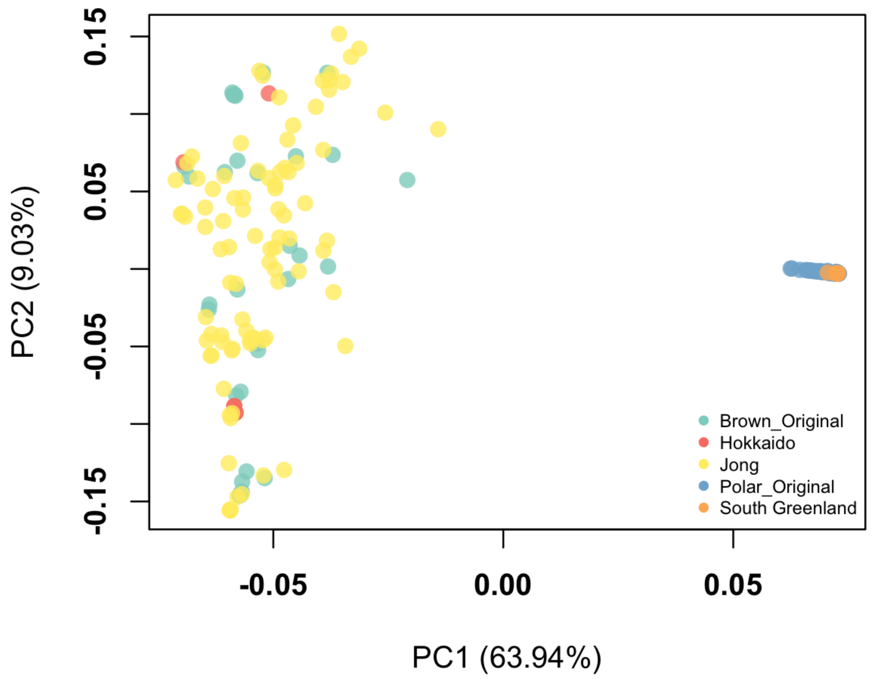


**Figure S3**. Principal component analysis of gene ABCC6 and the 50kb flanking regions using all modern individuals included in this study. 'Polar_Original' and 'Brown_Original' refer to the samples from Castruita, Westbury, and Lorenzen 2020. 'South Greenland', 'Hokkaido', represent new polar bear datasets from Laidre et al and Endo et al respectively. 'Jong' shows the Holarctic brown bear dataset from de Jong et al.

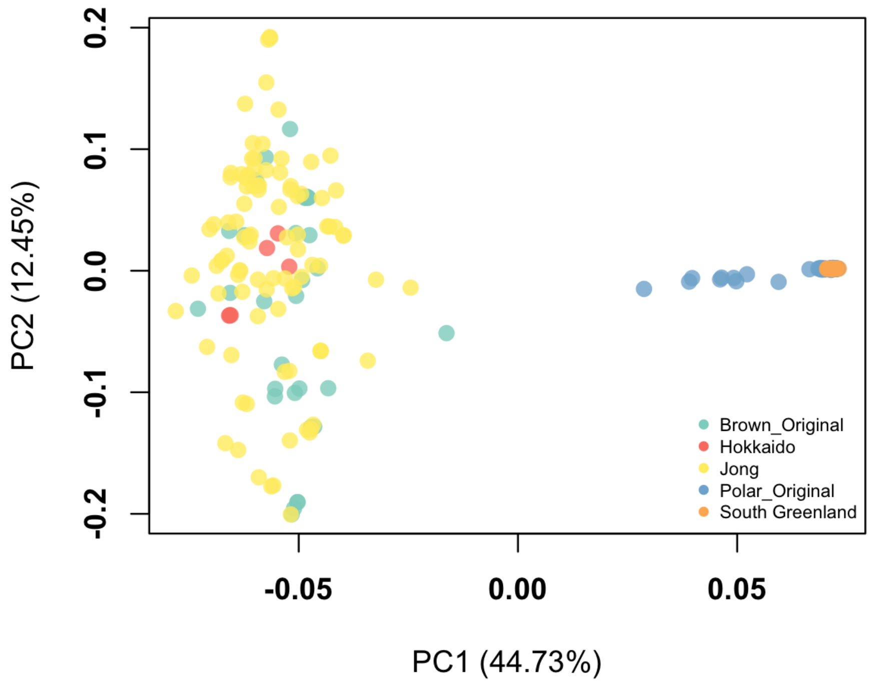


**Figure S4**. Phe principal component analysis of gene APOB and the 50kb flanking regions using all modern individuals included in this study. 'Polar_Original' and 'Brown_Original' refer to the samples from Castruita, Westbury, and Lorenzen 2020. 'South Greenland', 'Hokkaido', represent new polar bear datasets from Laidre et al and Endo et al respectively. 'Jong' shows the Holarctic brown bear dataset from de Jong et al.

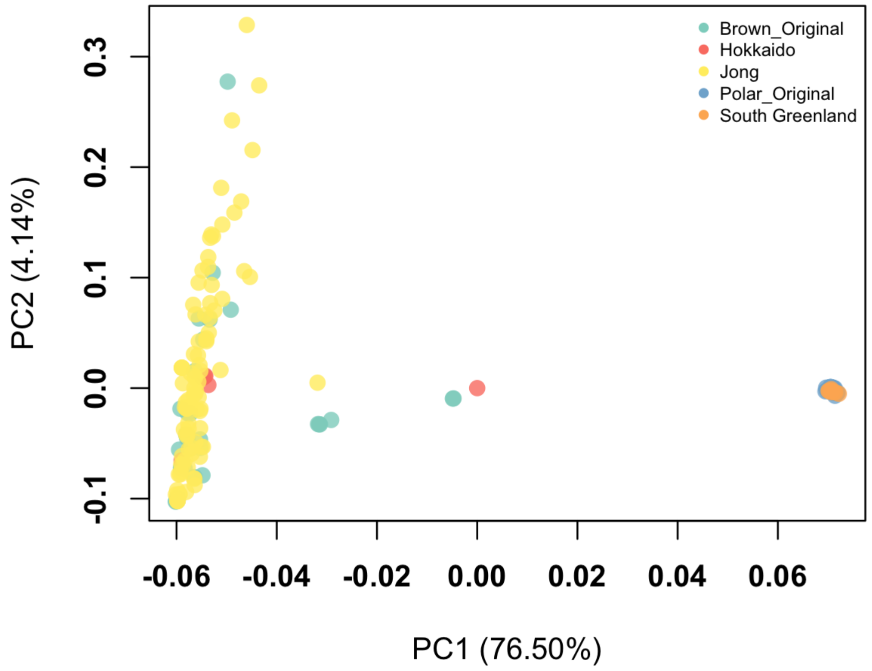


**Figure S5**. Principal component analysis of gene COL5A3 and the 50kb flanking regions using all modern individuals included in this study. 'Polar_Original' and 'Brown_Original' refer to the samples from Castruita, Westbury, and Lorenzen 2020. 'South Greenland', 'Hokkaido', represent new polar bear datasets from Laidre et al and Endo et al respectively. 'Jong' shows the Holarctic brown bear dataset from de Jong et al.

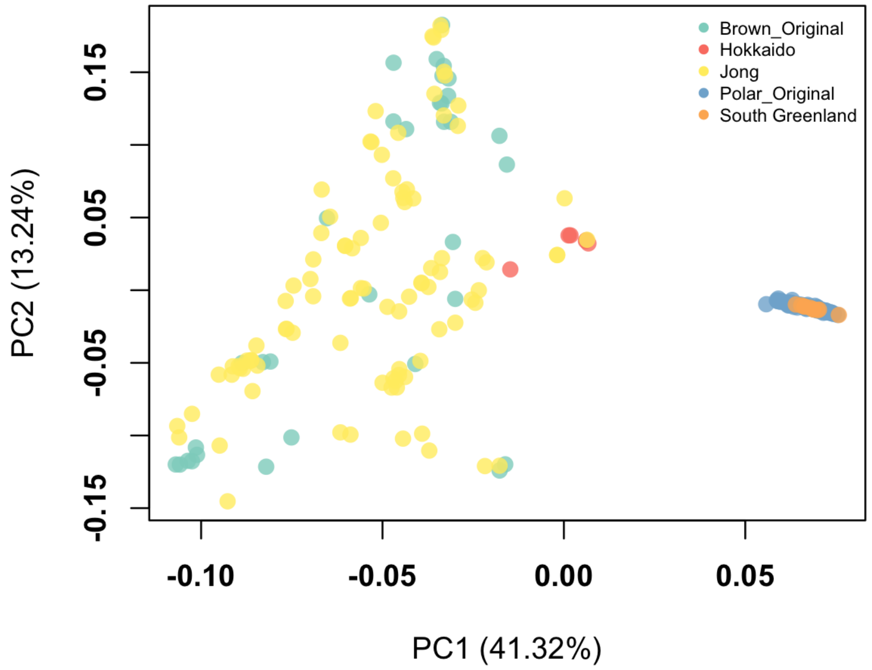


**Figure S6**. Principal component analysis of gene CUL7 and the 50kb flanking regions using all modern individuals included in this study. ‘Polar_Original’ and ‘Brown_Original’ refer to the samples from Castruita, Westbury, and Lorenzen 2020. ‘South Greenland’, ‘Hokkaido’, represent new polar bear datasets from Laidre et al and Endo et al respectively. ‘Jong’ shows the Holarctic brown bear dataset from de Jong et al.

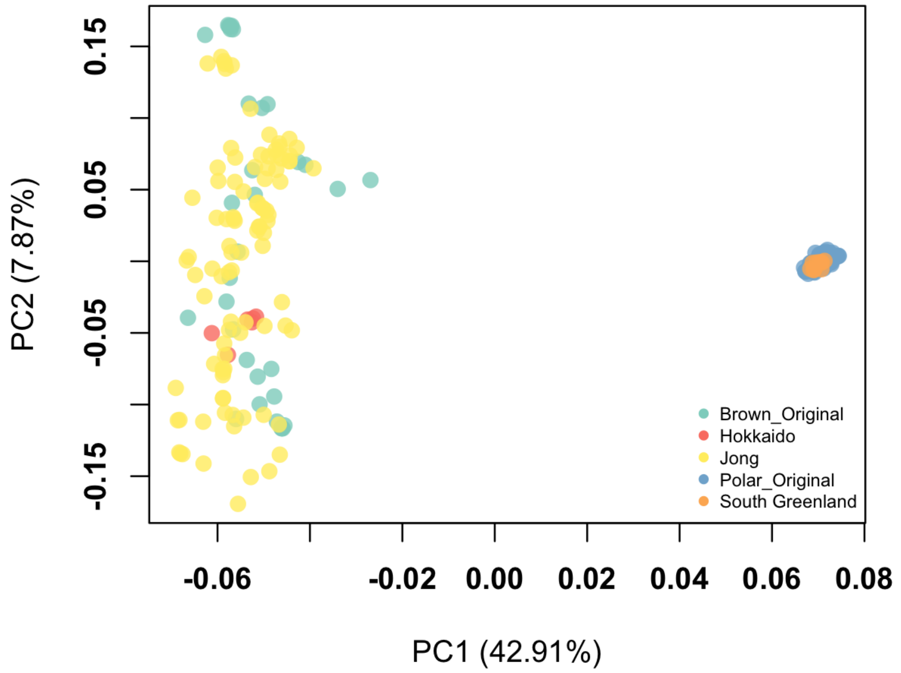


**Figure S7.** Principal component analysis of gene FCGBP and the 50kb flanking regions using all modern individuals included in this study. 'Polar_Original' and 'Brown_Original' refer to the samples from Castruita, Westbury, and Lorenzen 2020. 'South Greenland', 'Hokkaido', represent new polar bear datasets from Laidre et al and Endo et al respectively. 'Jong' shows the Holarctic brown bear dataset from de Jong et al.

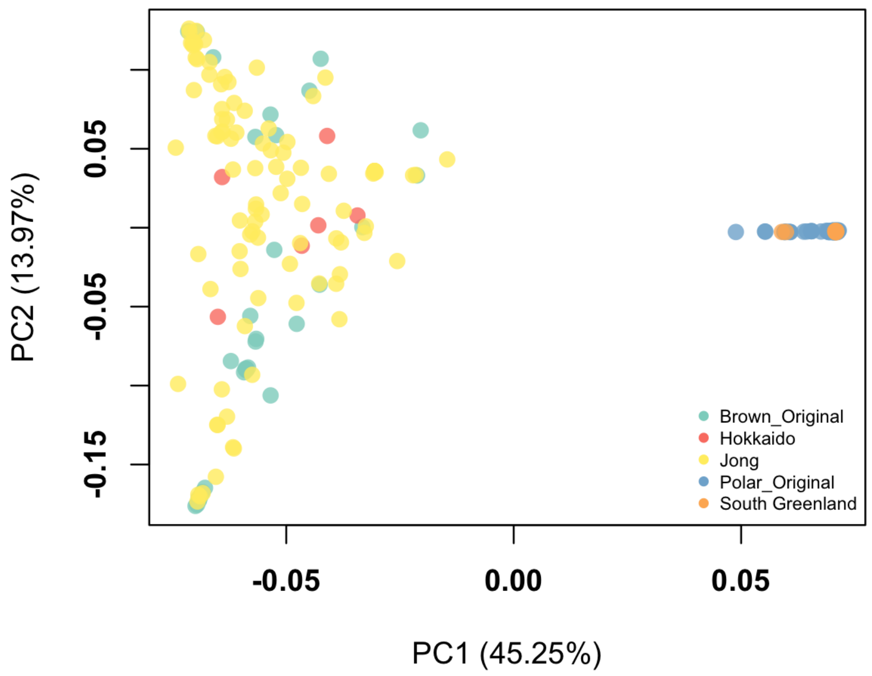


**Figure S8**. Principal component analysis of gene LYST and the 50kb flanking regions using all modern individuals included in this study. 'Polar_Original' and 'Brown_Original' refer to the samples from Castruita, Westbury, and Lorenzen 2020. 'South Greenland', 'Hokkaido', represent new polar bear datasets from Laidre et al and Endo et al respectively. 'Jong' shows the Holarctic brown bear dataset from de Jong et al.

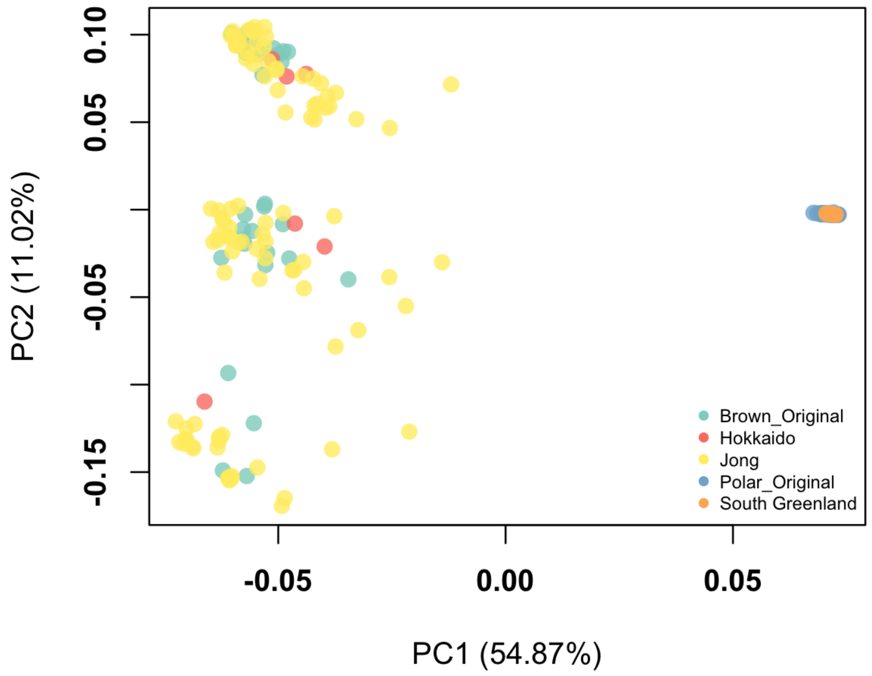


**Figure S9**. Principal component analysis of gene LAMC3 and the 50kb flanking regions using all modern individuals included in this study. 'Polar_Original' and 'Brown_Original' refer to the samples from Castruita, Westbury, and Lorenzen 2020. 'South Greenland', 'Hokkaido', represent new polar bear datasets from Laidre et al and Endo et al respectively. 'Jong' shows the Holarctic brown bear dataset from de Jong et al.

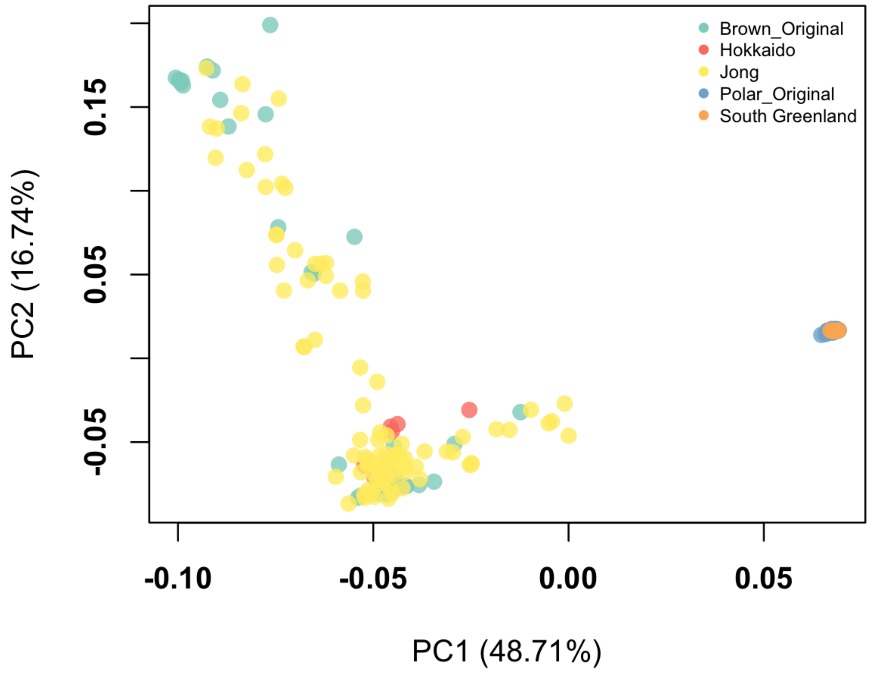


**Figure S10**. Principal component analysis of gene POLR1A and the 50kb flanking regions using all modern individuals included in this study. 'Polar_Original' and 'Brown_Original' refer to the samples from Castruita, Westbury, and Lorenzen 2020. 'South Greenland', 'Hokkaido', represent new polar bear datasets from Laidre et al and Endo et al respectively. 'Jong' shows the Holarctic brown bear dataset from de Jong et al.

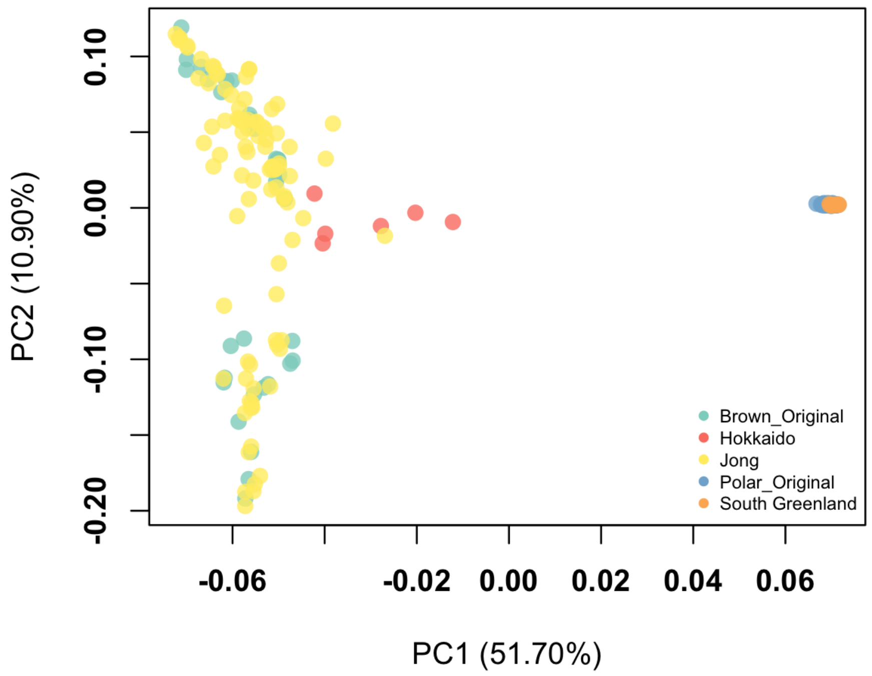


**Figure S11**. Principal component analysis of gene TTN and the 50kb flanking regions using all modern individuals included in this study. ‘Polar_Original’ and ‘Brown_Original’ refer to the samples from Castruita, Westbury, and Lorenzen 2020. ‘South Greenland’, ‘Hokkaido’, represent new polar bear datasets from Laidre et al and Endo et al respectively. ‘Jong’ shows the Holarctic brown bear dataset from de Jong et al.

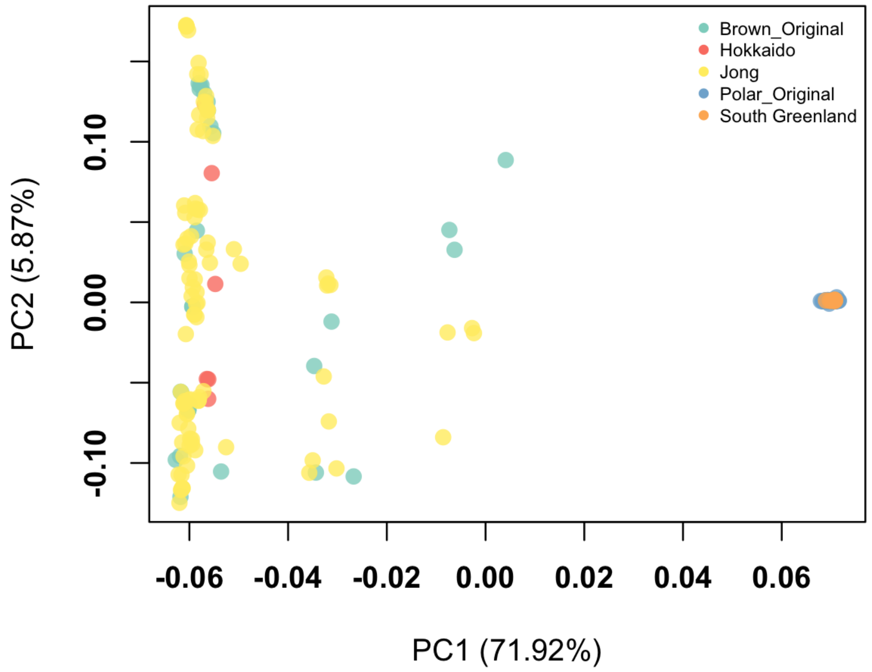


**Figure S12**. Principal component analysis of gene XIPR1 and the 50kb flanking regions using all modern individuals included in this study. 'Polar_Original' and 'Brown_Original' refer to the samples from Castruita, Westbury, and Lorenzen 2020. 'South Greenland', 'Hokkaido', represent new polar bear datasets from Laidre et al and Endo et al respectively. 'Jong' shows the Holarctic brown bear dataset from de Jong et al.
